# Digital Twin Approaches for Interpretable Side Effect Prediction in Drug Discovery

**DOI:** 10.1101/2025.10.14.682276

**Authors:** András Ecker, Gergely Szabó, János Szalma, Erzsébet Fichó, István Reguly, Attila Csikász-Nagy

**Affiliations:** Cytocast Hungary Kft, Budapest, Hungary; Faculty of Information Technology and Bionics, Pázmány Péter Catholic University, Budapest, Hungary

**Keywords:** drug discovery, side effects, machine learning, feature representation, interpretability, actionable insight

## Abstract

Artificial intelligence plays an ever-greater role in preclinical drug development, ranging from target identification and molecule design to ADME-Tox prediction; however, predicting side effects before performing clinical trials is still lagging behind. The best performing side effect predictors in the literature use either ATC codes, which are expert-derived features not even available at early stages, or graph neural networks based on chemical similarity, which - although use readily available features - are “black boxes” that do not deliver actionable insights. We argue that a paradigm shift is needed. Instead of using the latest neural network architectures that have proved worthy in other domains with a plethora of available data, one could use the off-targets of the compounds and build simple and interpretable predictors of side effects. To add another layer of biological realism, intricate biophysical mechanisms within the cells could also be simulated and used as features for training. Although not outperforming the current methods by a great margin, this digital twin-based model has the benefit of being interpretable, i.e., it puts biology behind the predictions. We showcase, with real-world examples, how the side effects predicted by this model can be interpreted and traced back to off-target proteins, and the complexes and signaling pathways in which they partake. In this way, the proposed model not only provides actionable insights, but in the future, may contribute to the amendment of secondary pharmacology assays.

**Highlights:**

- No standard tool is available to predict side effects in early-phase drug discovery.
- As publicly available side effect data is scarce, only simple models should be trained.
- Simple models trained on biorealistic features, such as off-target proteins, are interpretable.
- Interpreting predictions can highlight critical off-targets and are therefore actionable.

## 1 Introduction

Drug development is an intricate and risky process, often stretching over a decade and costing billions of dollars. Once efficacy is confirmed, the main cause for terminating clinical trials or even withdrawing drugs from the market is side effects. To alleviate the burden, several *in silico* methods have been established to predict potential side effects of drug candidates.

In the 2010’s classical machine learning methods, such as logistic regression, support vector machines (SVMs), decision trees, matrix factorization, knowledge graph-, and network-based methods dominated the field ^1,2^. These models combine features in a way that is easy to grasp and trivially invertible in the mathematical sense; thus the way their predictions are made can be interpreted. The features resulting in the best-scoring models include drug phenotype, e.g. the Anatomical Therapeutic Chemical (ATC) code, and drug chemical features, e.g., Tanimoto similarity of Morgan Fingerprints ^1^. While maximizing performance is important in the machine learning community, and the interpretability of these algorithms is a huge plus, the interpretation of these features is not really actionable by practitioners. Predicting side effects that resemble those of previously approved drugs that belong to the same ATC category according to the WHO, and backing it by chemical similarity, is just not pragmatic.

When the predictive performance of these models plateaued in the 2020’s, the focus shifted towards neural networks^3^, which gained traction in other biology-related domains, such as predicting the 3D structure of proteins ^4^. However, to achieve their superior performance, deep neural networks require an immense amount of training data, which in the case of adverse drug reactions (ADRs), is not publicly available. Furthermore, while neural networks build their own feature representation and mitigate the problem of feature engineering, without a thorough analysis of this representation, their predictions are much more difficult to interpret. This has earned them a *“black box”* designation and the distrust of non-experts (Figure 1C).

**Figure 1:**
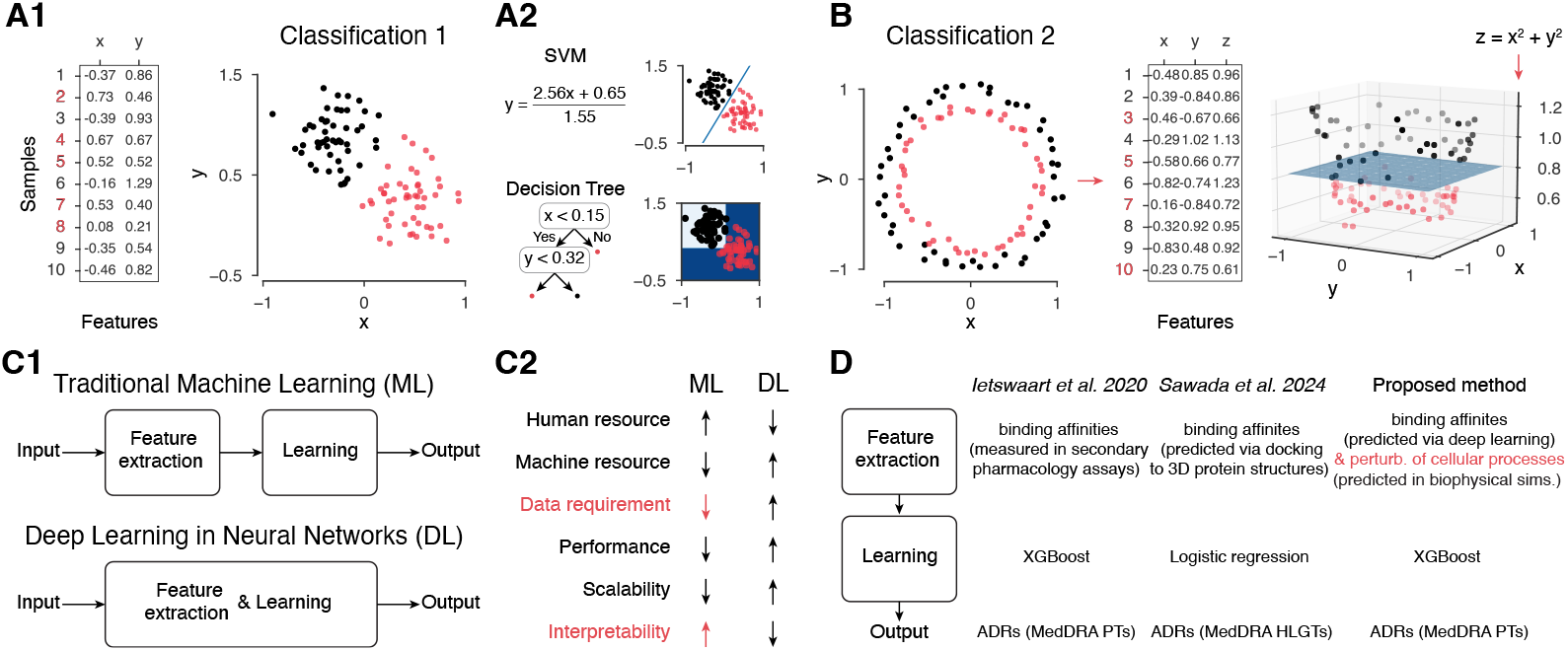
Overview of feature representation and machine learning models. **A1:** First example of a 2D classification task (black vs. red). Table on the left shows 10% of the data points in detail. Colors on the y-axis match the class of the given data point. **A2:** Two possible solutions of the task using the original features with either SVM (on top) or a decision tree (on the bottom). **B:** Second example of a 2D classification task, with a newly engineered feature (*z* as the sum of squares) which makes it solvable by an SVM in 3D. **C:** Overview of simple models using feature engineering and neural networks that build their own feature representation. **D:** Comparison of key elements of two recent models with our proposed one.

Thus, with the exception of a few notable examples discussed in the followings ^5–7^, early-stage drug development to date does not obtain actionable insights from available side effect prediction tools. In this Perspective, we argue that advancing the field requires training simpler models that leverage better features.

## 2 Simpler models using better features

### 2.1 Good feature representation

We will introduce the concept of good features with 2D classification tasks. When points belonging to two classes are detached blobs as the red and black dots in Figure 1A1, they can be easily separated by a line using their *x* and *y* coordinates as features. This can be done by using the two features simultaneously by a linear classifier, or by a more advanced SVM that maximizes the margins from the closest points (Figure 1A2 top). Another possible solution would be a decision tree, that builds tree-like structures using if-then rules considering only a single feature at a time (Figure 1A2 bottom). Real-world examples, however, tend to be more complicated and not easily separable using only the original input features. We illustrate this by classifying concentric circles (Figure 1B left), which cannot be solved in 2D by an SVM. A large decision tree would achieve better performance, but would overfit and be hard to interpret because of the high number of if-then decisions. On the other hand, by introducing a new feature as the sum of squares, resulting in a paraboloid in 3D would make the problem again easily solvable (Figure 1B right, blue hyperplane fitted by an SVM). This type of feature engineering is however time consuming and requires human intuition (Figure 1C2), which led to the rapid rise of employing neural networks that build their own feature representation while solving the given task (Figure 1C1). When these networks were small, it was feasible to understand the features learned^8^. However, since their parameters exceeded the billions, there is no straightforward way to interpret them. In computer vision applications, it is a common practice to verify whether the layers of the network behave similarly to brain areas of the visual stream processing the same information^9^. In other fields like systems biology, however, this is simply not possible.

As for coming up with good feature representations, one can still turn to basic pharmacology. Notably, drug candidate compounds are designed to target specific proteins and exert their effects on these *on-targets*. The landmark study of Campillos et al. ^10^ showed, that on-targets can be predicted from side effects. While they did it for drug repositioning, it inspired some of the early side effect prediction models to use on-targets as features^2^. On the other hand, while even on-target perturbation can lead to undesired effects, most ADRs arise from unintentional binding to other, *off-target* proteins. Secondary pharmacology assays are designed to test a few hundred important proteins linked to known side effects ^11^, thus if available, these serve as good features. Indeed, the seminal work of Ietswaart et al. ^6^ used a plethora of these measurements to build XGBoost models (ensembles of decision trees) predicting ADRs. Since laboratory measurements are only available in a few cases, Sawada et al. ^7^ used *predicted binding affinities* against the human proteome (Figure 1D). They then predicted side effects with a simple logistic regression from these, which achieved higher accuracies than the same model trained on the same labels with traditional features. After clustering the predicted binding affinity matrix, they found that ATC codes were overrepresented in some of their clusters ^7^. Similar conclusions were reached by Galeano et al. ^5^, who used matrix decomposition to build low dimensional representations of both drugs and side effects. Some of their 10*D* drug feature vectors were also significantly correlated with ATC codes. In summary, correlation with ATC codes that contain valuable, expert-derived information about the drug can be seen as a good sign of well derived features. Note however, that ATC codes themselves are only determined in the late phase of drug development (usually after clinical trials), thus they cannot be used to predict potential ADRs in the early phases of the pipeline.

For even more features, one can turn again to systems biology, and acknowledge the fact that the target proteins partake in a network of protein-protein interactions (PPIs), form protein complexes, and participate in signaling pathways. While powerful and widely used ^1^, simple graph-based approaches on PPI networks neglect the fact that both protein abundances and binding sites are limited. To take this into account and study complex formation under constrained resources, we built an agent-based simulator that mimics the behavior of proteins within the cell ^12^; the CYTOCAST DIGITAL TWIN Cell^™^ (Figure 2A top). Relevant data was sourced from several databases: protein abundances from PaxDB^13^, protein domains from Pfam ^14^, PPIs from STRING^15^, domain-domain interactions (DDIs) from 3did ^16^, and reference complexes from ComplePortal ^17^ and the Reactome Knowledgebase ^18^. The simulator also tracks PPIs of signaling pathways, parsed from the Reactome Knowledgebase ^18^. Furthermore, in parallel with Sawada et al. ^7^, we developed our own binding affinity predictor (detailed in a future manuscript) to predict targets of drug compounds from a list of 7390 druggable proteins (Figure 2A-B). Using these predicted targets we were able to extend the above simulations by adding interactions between the drug compound and domains of the predicted targets, which perturb the complexome and the signaling pathways (Figure 2A bottom). These significantly perturbed complexes and pathways are highly preprocessed bio-realistic features, which can also be used for side effect prediction. To derive them for a new compound of interest, only its structure is required, thus they are readily applicable in the earliest stage of drug development.

**Figure 2:**
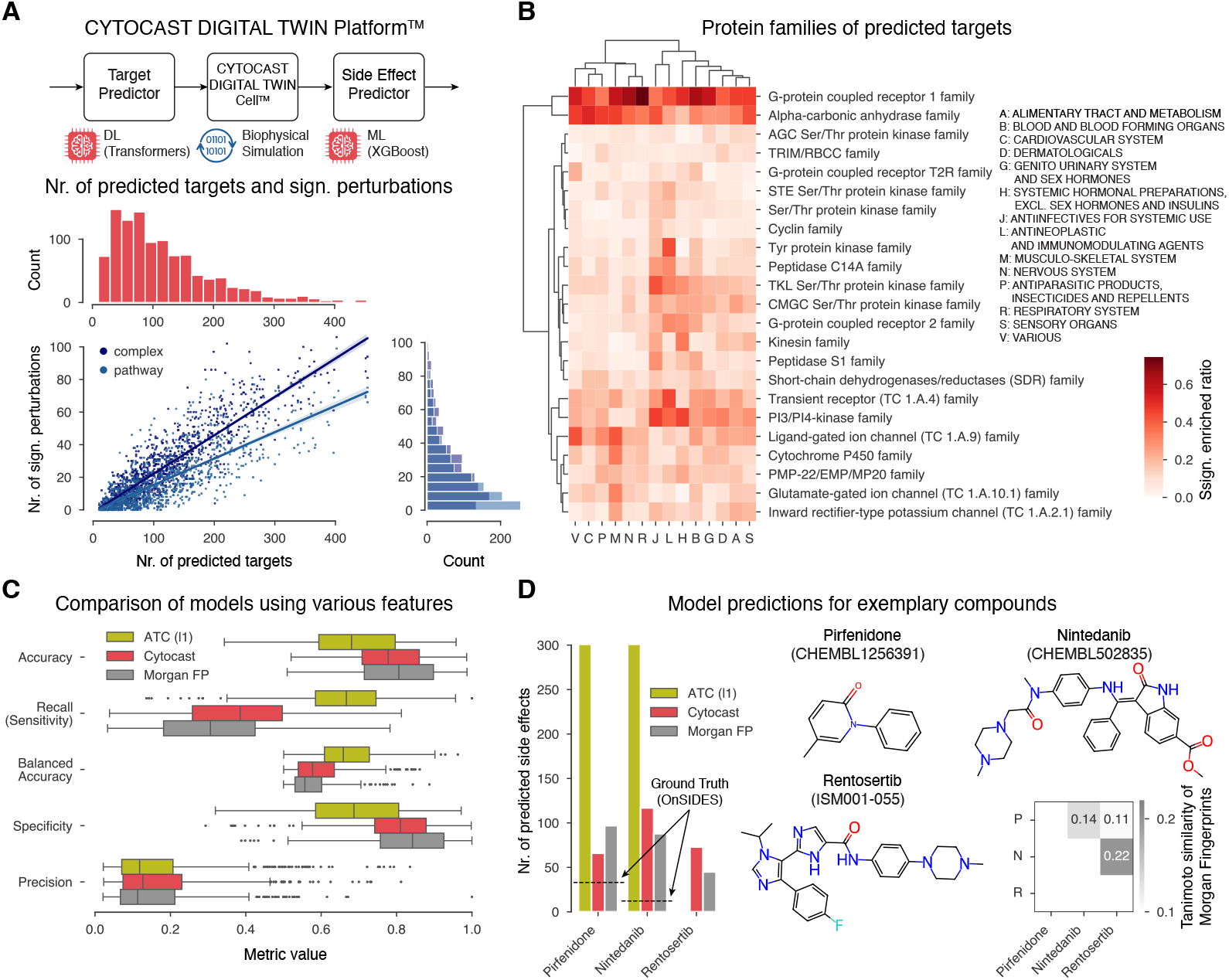
Models trained on various features. **A:** Overview of Cytocast’s pipeline on top, and simulation derived features on the bottom: number of significantly perturbed complexes and pathways (775 and 206 unique ones respectively) as a function of the number of predicted targets (4150 unique off-targets for 1069 training drugs). **B:** Enrichment of protein families of predicted targets. Heatmap shows the ratio of drugs from the ATC category for which the given family was significantly enriched (Fisher’s exact test, with significance threshold of 0.05). Only the most common families are shown. **C:** Summary of various metrics of three different side effect prediction models (same training labels, same models, but different features) **D:** Comparison of model predictions for selected compounds. Number of predicted side effects on the left, structure of compounds (visualized using RDKit), and similarity of the compounds, on the bottom right. On the left the ground truth side effect count as well as the ATC (level 1) code-based prediction for Renstosertib are missing, because it is in active development.

### 2.2 Training simple models

In this section, we present results obtained by training XGBoost models with the same codebase on the same side effect data (59’803 drug - side effect pairs from OnSIDES ^19^, with 1069 unique drugs and 561 unique side effects as MedDRA ^20^ Preferred Terms). The purpose of this exercise is to showcase how the feature representations introduced above compare with traditional features like ATC codes and drug fingerprints in terms of performance. Due to the imbalanced occurrence of side effects (with only a few drugs causing specific side effects), we trained separate models for each side effect to balance drugs across the cross-validation folds. While training separate models is common practice in the field ^2,3,6,7^, using balanced accuracy - defined as the mean of sensitivity and specificity - as the primary evaluation metric to address the imbalance is new. The models trained on ATC level 1 codes are superior in sensitivity (also known as recall), while the ones on Morgan fingerprints in specificity (Figure 2C). The models trained on our newly proposed features (labeled as Cytocast on Figure 2C) balance these two, and outperform the others in terms of precision.

The models were further challenged in a real world setup, using compounds, that were not part of the training set. Idiopathic pulmonary fibrosis (IPF) is a progressive, chronic interstitial lung disease of unknown etiology and has a median survival prognosis of 3-5 years, currently treated with Nintedanib and Pirfenidone ^21^ (Figure 2D). Both clinical trials and pharmacovigilance data mining highlighted slightly different side effect profiles of the two drugs^21,22^. As they share the same ATC level 1 code (L), the model trained on those cannot differentiate between them and predicts the same 301 side effects for both (Figure 3E left). Furthermore, according to the OnSIDES database ^19^ there are only 12 and 30 ADRs associated with the compounds (Figure 2D left, dashed lines). Accordingly, the ATC-based model vastly overpredicts, which also explains its superior recall (also known as true positive rate). Albeit both compounds are effective in slowing down disease progression, neither can reverse its course, thus finding new compounds to treat IPF is an active area of research. The third exemplar compound, Rentosertib, is an AI-discovered IPF medication in phase 2 clinical trial ^23^ (Figure 2D). Since it does not yet have an ATC code, no predictions can be made with this model (Figure 2D left). While the problems could be mitigated by predicting even the ATC code and adding more levels to the feature set, it illustrates the problems of learning on not readily available features and predicting ADRs that are general representatives of the given drug category, and not specific to the compound itself. Concurrently, the Morgan fingerprint-based models can be used in the preclinical phase and are able to differentiate between the compounds, the interpretation of their predictions would only highlight similarities in the structures themselves (Figure 2D bottom right). Furthermore, they not only fall short of the ones using biorealistic features in balanced accuracy of the side effects (0.57 ± 0.06 vs. 0.6 ± 0.08, see Figure 2C), but also that of the drugs (0.58 vs. 0.65, and 0.48 vs. 0.6 for Pirfenidone and Nintedanib respectively).

**Figure 3:**
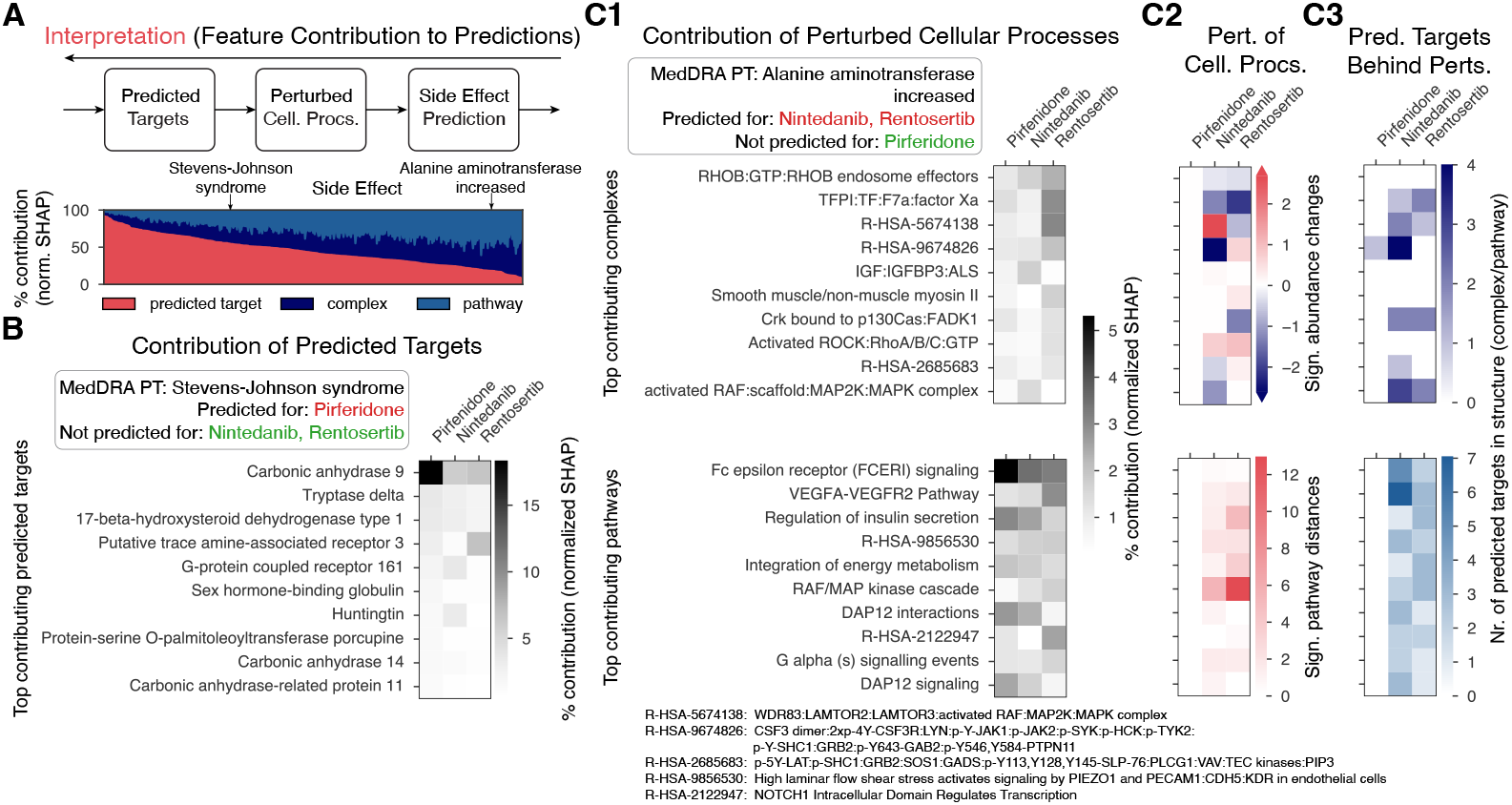
Interpretability of predictions. **A:** Flowchart illustrating working backwards when interpreting the (Cytocast) model’s predictions (see C for details) on top. Average contribution (normalized SHAP values) of the three different feature types in the training data on the bottom. Side effects (on the x-axis) are sorted by the contribution of predicted targets. Arrows indicate the two side effects shown in detail on B-C. **B:** First example of interpretability for a selected - predicted target dominated - side effect. The heatmap shows the contribution of the top ten predicted targets (selected based on, and order by Pirfenidone). **C:** Second example of interpretability for a selected - perturbed complex and pathway dominated - side effect. **C1:** The heatmaps show the contribution of the top ten complexes on top, and pathways on the bottom (selected based on, and order by the sum of Nintedanib and Rentosertib). **C2:** The heatmap on top shows the significantly (Mann–Whitney U test, with significance threshold of 0.001) perturbed complexes in the simulation. Positive/red hues indicate an increase, negative/blue hues a decrease in complex abundance, while white squares represent non-significant changes. The heatmap on the bottom shows the significantly perturbed signaling pathways in the simulation. Pathway distances are calculated as Euclidean distances of PPI count vectors. Significance test and missing values as for complexes above. **C3:** The heatmaps show the number of predicted targets, that are part of the complexes on top and pathways on the bottom.

### 2.3 Interpreting predictions

In this section we highlight the main strength of the proposed Cytocast model and show how actionable insight can be gained from interpreting its predictions. From the various available mathematical frameworks, we have chosen the SHapley Additive exPlanations (SHAP) analysis, which distributes contribution among the features ^24^, thus helping to identify which off-target proteins, perturbed complexes and pathways lead to an ADR being predicted. We observed that the contribution of the feature types in our training set varies across side effects (Figure 3A).

For the first example, we selected “Stevens-Johnson syndrome”, an off-target dominated side effect, which we correctly predicted for Pirfenidone. By looking into the SHAP values of the potential off-target proteins, we found that the prediction was 18% based on Carbonic anhydrase 9 (Figure 3B) which was indeed a predicted off-target of Pirfenidone. The same off-target was also the top contributor (8%) to a related true positive prediction, toxic epidermal necrolysis. It is important to note here that this analysis does not necessarily describe the biological mechanisms behind the ADRs, rather just showcases how our model understands them. However, in the case of carbonic anhydrase inhibitors there is a known link to severe skin-related side effects such as the two above ^25^.

For our second example, we selected “alanine aminotransferase increased”, a side effect characterized by perturbations in complexes and pathways. This adverse event was also reported among the four most frequently occurring ADRs in the phase 2a clinical trial of Rentosertib by Xu et al. ^23^. Perturbations in the VEGFA-VEGFR2 signaling pathway are known to be associated with increased alanine aminotransferase levels ^26^, while some of the remaining pathways and complexes - especially those linked to insulin secretion and the MAPK pathway - are also intertwined with it (Figure 3C1). Most, but not all of these were perturbed in the simulations for Rentosertib (and Nintedanib for which we also predicted it), which point out that features with value 0 can also heavily influence the predictions (Figure 3C2). These perturbed complexes and pathways can ultimately be linked back to off-target proteins (Figure 3C3). Elimination of these off-targets, or even just lowering the compounds binding affinity to them could be an actionable step against the ADR, or if that is not possible it could help with patient stratification when preparing for the clinical trial. Furthermore, due to the fully *in silico* nature of the study these hypotheses about off-target reduction can be tested within the same framework.

## 3 Discussion

In summary, while *in silico* tools are already in use for ADME-Tox predictions^27^ and digital twin technology is revolutionizing healthcare in general^28^, there is no industry standard tool that predicts ADRs. Some of the methods available in the literature appear like to focus primarily on maximizing performance ^29^, without addressing neither practitioner needs nor the availability of the data in inference time. The bigger problem is that of limited data. The most used side effect resource to date is SIDER^30^, which due to the lack of funding, was last updated more than 10 years ago. Many first-in-class drugs appear every year, and not taking these into account seriously limits the domain coverage of the trained models. Pharmacovigilance data, e.g., deposited to FAERS^31^ is also a valuable source of information. This applies even to drugs that have been on the market for a long time. For these reasons, we trained our models on side effects extracted by large language models (LLMs) from the latest drug labels and stored in the OnSIDES database ^19^. While being up to date, the LLM-based extraction also has its own error rate, and training models on labels that are not only limited and sparse, but also noisy is a hard problem. Care must be taken in regularizing the models and correctly evaluating their performance in stratified cross-validation loops. Furthermore, although the most useful predictions would be about severe side effects that could put an end to clinical trials or lead to the withdrawal of marketed drugs, there is as of yet no publicly available data about ADRs that terminated clinical trials.

Acknowledging the paucity of available data, we strongly argue against training complex models, which while being able to extract highly entangled relationships, need to see large amounts of samples to do so. The added benefit of traditional machine learning techniques is that they are mathematically invertible or if not, at least interpretable. To perform well, however, these models need good features, like measured ^6^ or predicted ^1^ off-target proteins. Our contribution lies in proposing to use even more of these biologically relevant features. Although one might argue that the proposed digital twin cell-based features add another layer of complexity to the model, we would highlight that this feature representation is at least data-driven, not performance-driven and can be traced back to off-targets. This unique aspect of the proposed method may contribute to the extension or revision of secondary pharmacology assays, routinely run in drug discovery settings ^11^. Due to its non-black box nature, this technology already provides actionable insights, and in the future it might even aid the drug design process.

## Acknowledgements

The authors thank all employees of Cytocast Hungary Kft., especially advisor György M. Keserű and commercial development leader Paul Lebeau for their helpful comments on earlier versions of the manuscript.

